# Genetic mapping of a spontaneous short-grain mutation reveals a novel loss-of-function allele of *SRS3* in rice

**DOI:** 10.64898/2026.08.03.742661

**Authors:** Maria Montiel, Brijesh Angira, Jonathan K. Richards, Adam Famoso

## Abstract

Spontaneous mutations are a rare but important source of novel genetic variation, yet their detection and characterization within active breeding programs are seldom documented at gene-level resolution. Grain size and shape are key determinants of rice quality, yield, and market classification. Here, we report the discovery and genetic characterization of a spontaneous short-grain (SG) mutation arising in the long-grain wild-type (WT) advanced breeding line RU2002174 from the LSU AgCenter Rice Breeding Program. The SG phenotype was first observed in 2019 and segregated in subsequent generations as a single recessive gene across both *indica* and *japonica* genetic backgrounds. Genetic mapping localized the mutation to a 41.6 kb interval on chromosome 5. Whole-genome sequencing identified a single candidate causal variant: a G→T transversion in exon 4 of *SRS3* (Os05g06280), introducing a premature stop codon and resulting in a truncated protein.

This allele was absent from representative U.S. breeding germplasm and the IRRI 3K SNP database, demonstrating that it represents a novel spontaneous loss-of-function allele of a previously characterized grain-size gene. These findings document the real-time emergence of functional genetic variation in elite rice germplasm and highlight the importance of monitoring off-types during seed increase and purification in breeding programs. They also provide additional insight into the role of kinesin-mediated cell elongation in determining rice grain architecture.

## 1. Introduction

Rice (*Oryza sativa* L.) is a staple food for more than half of the world’s population and plays a central role in global food security [1]. As demand continues to increase, rice breeding programs must simultaneously improve grain yield and quality [2, 3, 4]. Grain size and shape are key determinants of rice appearance quality, influencing grain weight, milling recovery, yield, and consumer acceptability [5, 6, 7, 8].

In the United States, rice varieties are classified by milled kernel length as long, medium, or short grain and by length-to-width ratio as bold, medium, or slender [9, 10, 11]. These classifications are tightly linked to market channels and consumer preferences. Grain size and shape are under strong genetic control, and extensive genetic studies and mutant resources have facilitated the characterization of grain morphology in rice [12, 13]. Genetic studies have identified numerous loci influencing grain morphology, including *GS3* [13], *GL3.1* [14, 15], *An1* [16], *GLW7* [17], *GS2* [18], qGL7.1 [19], *GW2* [20], *GW3.1* [21], *GW5* [22], *GS5* [23], and *GIF1* [24]. Natural variation and human selection have shaped this diversity during rice domestication and breeding [25].

In addition to natural allelic diversity, induced mutagenesis has been widely used to identify genes controlling grain size and shape. Several grain-shape mutants have been described from irradiation- and chemically induced populations. The *lgs1* mutant, derived from γ-Co60 irradiation, is a semi-dominant gene on chromosome 2 that increases grain size and enhances cold tolerance [26]. The *gs9–1* mutant is semi-dominant and affects gibberellic acid biosynthesis, resulting in reduced cell length and number [27]. The EMS-derived *smg11* mutant influences grain and panicle size through altered cell expansion [28]. The *SMALL AND ROUND SEED 3* (*SRS3*) gene was originally identified in a N-methyl-N-nitrosourea-induced population and shown to encode a kinesin-13 protein that regulates cell elongation during seed development, producing shortened and rounder seeds [29, 30].

Although induced mutants have contributed substantially to the identification of grain-size genes, spontaneous mutations represent a continuous source of novel genetic variation in crop plants. While spontaneous mutation rates in higher plants are low, such mutations arise repeatedly and can persist within breeding populations [31, 32, 33, 34]. From a breeding perspective, spontaneous mutations are often undesirable because they can compromise varietal uniformity, stability, and market classification. Despite their practical importance, spontaneous mutations arising within advanced breeding lines are rarely documented from field observation through genetic mapping and molecular identification.

Here, we describe the discovery and characterization of a spontaneous short-grain mutation that arose in the long-grain breeding line RU2002174 from the LSU Rice Breeding Program. The objectives of this study were to confirm that the phenotype resulted from a spontaneous mutation rather than seed contamination, determine its inheritance across genetic backgrounds, and map and identify the underlying gene and candidate causal nucleotide change. By tracing a spontaneous off-type from field observation to gene-level resolution, this study documents the continued emergence of functional genetic variation in advanced breeding materials and provides additional insight into the genetic control of rice grain architecture.

## 2. Materials and Methods

### 2.1. Plant Material

A spontaneous mutant exhibiting a distinct short-grain (SG) phenotype was identified in 2019 within the LSU AgCenter Rice Breeding Program at the H. Rouse Caffey Rice Research Station (HRCRRS) near Crowley, Louisiana. The SG phenotype was first observed as an off-type in seed-increase panicle rows of the long-grain (WT) experimental breeding line RU2002174. During purification, SG plants were rogued, and hand-selected WT panicles were advanced to the 2019–2020 winter nursery in Lajas, Puerto Rico. Field management followed standard practices described in the Louisiana Rice Production Handbook [35].

Segregation of SG within Puerto Rico panicle rows indicated genetic heterogeneity within RU2002174. From a segregating row, both SG and WT panicles were selected and advanced. The RU2002174 SG mutant was crossed to the WT long-grain *indica* line IC206 to generate a mapping population of 1,031 F₂ individuals. A second mapping population consisting of 100 F₂ plants was developed by crossing the SG mutant to the long-grain tropical *japonica* cultivar ‘Avant’ [36].

For high-resolution fine-mapping, 14 recombinant F₂ plants from the IC206 × RU2002174 population were selected based on recombination within the target interval and self-pollinated to generate F₃ progeny-test families. A total of 251 F₃ individuals from these families were evaluated. To assess whether the SG allele was present in U.S. breeding germplasm, a Native Trait Panel (NTP) of 380 lines representing genetic diversity commonly used in U.S. rice breeding programs was also evaluated [37, 38, 39, 40].

### 2.2 Phenotyping Methods

Grain phenotype was scored by visual inspection of mature seed, with each plant classified as short grain (SG) or long grain wild type (WT) based on seed morphology. The distinction between SG and WT phenotypes was readily apparent visually and was supported by quantitative grain measurements (Supplementary Table S1).

Because the SG phenotype is recessive, F₃ phenotypic ratios enabled discrimination between homozygous WT and heterozygous F₂ genotypes, allowing refinement and localization of the SG locus among lines with this recombination pattern and recombination breakpoints within the mapped region.

### 2.3. Genotyping

Initial genotyping was performed on 147 F₂ individuals from the IC206 × RU2002174 population using the LSU500 SNP marker set [40]. LSU500 is an AmpSeq-based genotyping platform optimized for U.S. rice germplasm and implemented via Agriplex Genomics [40]. Of the 550 markers assayed, 312 SNPs were polymorphic with a minor allele frequency (MAF) > 0.2, with polymorphic marker counts ranging from 14 to 49 per chromosome (mean = 26).

Fine-mapping on chromosome 5 was conducted using PCR Allele Competitive Extension (PACE) assays [41, 42] on the LGC SNPline platform [43]. Genomic DNA was extracted using a HotSHOT protocol [44, 45]. Initial fine-mapping used eight existing PACE SNP markers to identify recombinants across the target region. Marker density was subsequently increased with 25 additional PACE assays designed using sequence comparisons between RU2002174 WT and IC206. Marker identifiers, physical positions, and primer sequences are provided in Supplementary Table S2.

In cases where direct genotyping of some F₂ individuals was not possible during fine-mapping as marker density increased, F₂ genotypes were inferred from marker allele segregation patterns observed in F₃ families.

#### 2.3.1. Sequencing

Whole-genome sequencing (WGS) was conducted on seven lines: WT RU2002174, WT IC206, the RU2002174-derived SG mutant, two F₂ individuals homozygous for the SG allele across the mapped region, and two F₂ individuals homozygous for the WT allele across the region.

Leaf tissue was collected and stored at −20 °C prior to DNA extraction. Samples were homogenized using a Bullet Blender with 0.5-mm and 2.0-mm zirconium beads. DNA extraction followed [46], and DNA quantity and integrity were assessed using a Qubit 3.0 fluorometer and agarose gel electrophoresis. Sequencing libraries were prepared using the IDT xGen DNA EZ Kit and sequenced on an Illumina NovaSeq X Plus platform to generate 150-bp paired-end reads, with a mean output of approximately 228.6 million read pairs per sample (∼90× coverage).

Read quality was assessed using FastQC [47]. Adapter sequences and low-quality bases were removed using Trimmomatic [48]. Trimmed reads were aligned to the *Oryza sativa* Nipponbare reference genome [49] using BWA-MEM [50]. Alignment files were converted to sorted and indexed BAM format using SAMtools [51]. Variant calling for SNPs and indels was performed using the Genome Analysis Toolkit (GATK) HaplotypeCaller, followed by joint genotyping using CombineGVCFs and GenotypeGVCFs [52]. Because all lines were fixed across the target region, heterozygous calls were set to missing. Variants were filtered using VCFtools with a minimum depth of 10 and genotype quality ≥ 30 [53]. All genome positions are based on IRGSP1 annotations.

### 2.4. Statistical Analysis

Single-marker analysis of variance was conducted using JMP [54] with genotype data from the LSU500 marker panel. Genotypes were quality-filtered using the *snpReady* package in R [55, 56], applying a minor allele frequency threshold > 0.2 and a minimum call rate of 0.95. Single-marker ANOVA was used to support coarse mapping of the SG locus.

Chi-square (χ²) tests were conducted in R using the chisq.test function to evaluate deviations from the expected 3:1 WT:SG segregation ratio for a single recessive gene [57].

## 3. Results

### 3.1. Observation of the Short Grain (SG) Mutant Line and Purification

A short-grain (SG) phenotype was first observed in 2019 in panicle-increase rows of the F4:F5 experimental breeding line RU2002174 at the H. Rouse Caffey Rice Research Station. RU2002174 is a long-grain tropical *japonica* breeding line from the LSU Rice Breeding Program (Fig. 1). To purify the line, visually normal long-grain (WT) panicles were hand-selected and advanced to the 2019–2020 winter nursery in Puerto Rico, where several panicle rows segregated at an approximate 3:1 WT:SG ratio.

**Fig 1:**
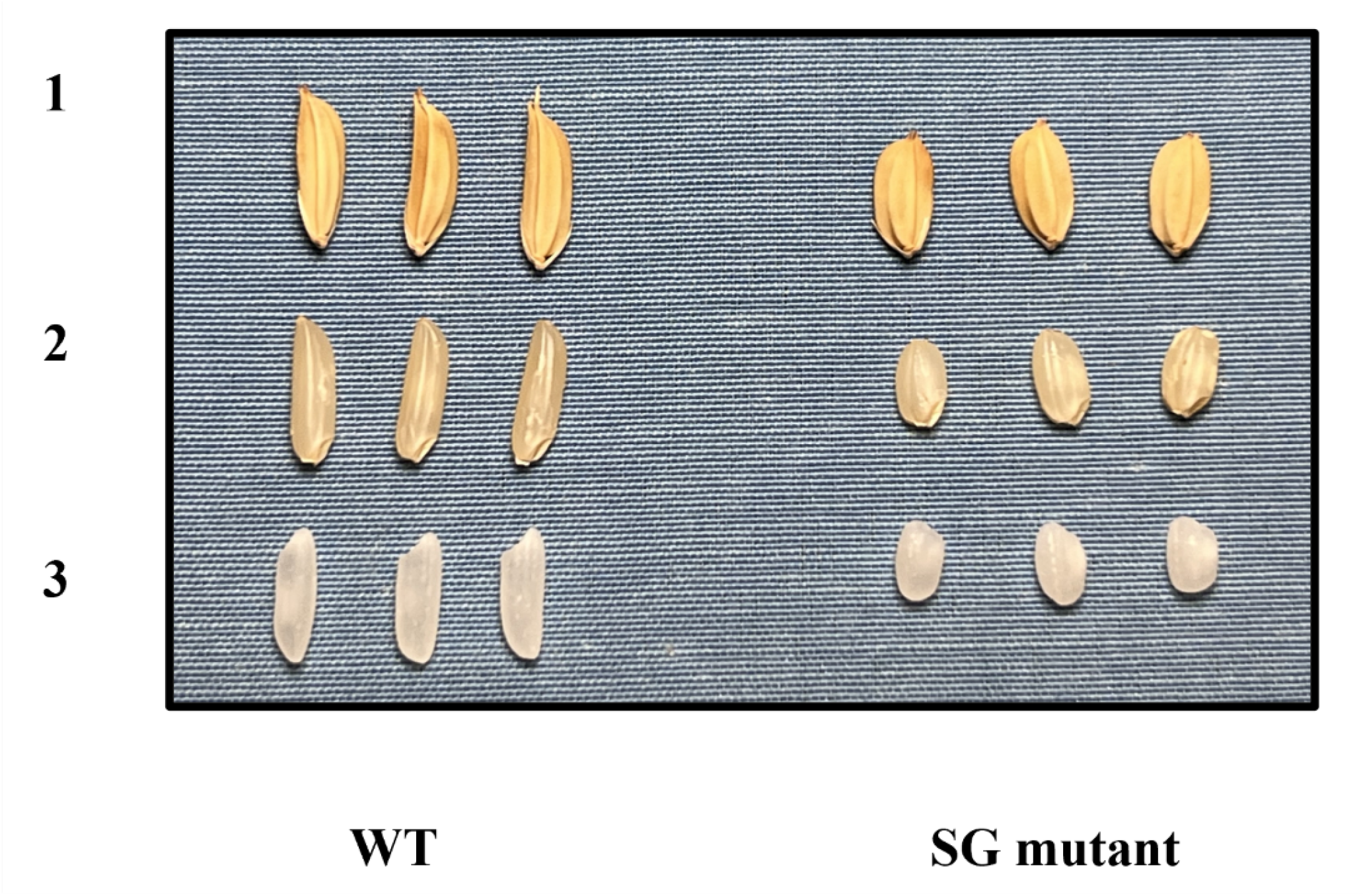
Grain morphology comparison between wild-type (WT) and short-grain (SG) phenotypes of RU2002174. Representative rough rice, brown rice, and milled rice grains are shown. Grain size differences (SG:WT ratios) were: rough rice L = 0.828, W = 1.280, T = 1.002; brown rice L = 0.633, W = 1.217, T = 0.937; and milled rice L = 0.607, W = 1.255, T = 1.026. Individual grain measurements are provided in Supplementary Table S1.

To determine whether the SG phenotype resulted from seed contamination or genetic mutation, SG and WT panicles from a single segregating row were genotyped using 550 genome-wide SNPs from the LSU500 marker set. SG and WT panicles were identical at 97% of SNP loci, indicating that the phenotype was unlikely to result from seed mix-up or outcrossing and instead reflected residual heterozygosity within the F4-derived line. Based on the absence of the SG phenotype in 2018 and its segregation in 2019, the mutation was inferred to have arisen during seed increase of RU2002174 in 2017, where it would have been present in the heterozygous state and phenotypically WT.

Grain morphology differed substantially between WT and SG plants. Compared with WT, the SG phenotype exhibited reductions in grain length ranging from approximately 17% in rough rice to nearly 40% in milled rice, while grain width increased by approximately 22–28%, resulting in a pronounced reduction in length-to-width ratio across rough, brown, and milled rice (Fig 1; Supplementary Table S1).

### 3.2. Phenotypic Inheritance and Rough Mapping of SG Phenotype in an F_2_ Population

To assess inheritance of the SG phenotype, the RU2002174 SG mutant was crossed to the long-grain *indica* line IC206 to generate an F₂ population. Among 147 F₂ individuals, 105 exhibited the WT phenotype and 42 exhibited the SG phenotype, consistent with a 3:1 segregation ratio for a single recessive gene (χ² = 0.31, *p* = 0.58). To confirm inheritance across genetic backgrounds, the SG mutant was also crossed to the long-grain tropical *japonica* cultivar ‘Avant’. Among 100 F₂ plants, 77 were WT and 23 were SG, again consistent with single-gene recessive inheritance (χ² = 0.21, *p* = 0.64).

For initial mapping, the 147 F₂ individuals from the IC206 cross were genotyped with the LSU500 SNP panel, of which 312 markers were polymorphic. Genome-wide single-marker analysis identified significant associations between the SG phenotype and markers on chromosome 5. Three SNPs located at 0.99, 4.13, and 4.59 Mbp showed strong association with the SG trait (Supplementary Table S3).

Haplotype analysis using markers at 0.99 and 4.13 Mbp indicated that the SG locus was located within this interval and confirmed recessive inheritance (Table 1). All parental genotype classes (WT:WT, WT:SG, SG:SG) were consistent with observed phenotypes. Recombinant individuals with homozygous WT alleles at one marker and heterozygous alleles at the other consistently exhibited the WT phenotype, whereas individuals carrying homozygous SG alleles at one marker showed both WT and SG phenotypes depending on the second locus. These results supported localization of the SG gene between 0.99 and 4.13 Mbp on chromosome 5.

**Table 1.** SNP haplotypes and genotype classes in the F₂ population showing phenotype counts per class. Allele classifications at each marker represent Homozygous long-grain parent allele (WT:WT), Homozygous short-grain mutant parent allele (SG:SG), and Heterozygous (WT:SG).

| Phenotype |  | Marker Position and Allele |  |
| --- | --- | --- | --- |
| Short Grain | Long Grain | 0.99 Mbp | 4.13 Mbp |
| 0 | 39 | WT:SG | WT:SG |
| 0 | 24 | WT:WT | WT:WT |
| 0 | 18 | WT:WT | WT:SG |
| 0 | 11 | WT:SG | WT:WT |
| 0 | 1 | SG:SG | WT:WT |
| 1 | 0 | WT:WT | SG:SG |
| 6 | 7 | SG:SG | WT:SG |
| 12 | 5 | WT:SG | SG:SG |
| 23 | 0 | SG:SG | SG:SG |
| Totals |  |  |  |
| **42** | **105** |  |  |

### 3.3 Fine-Mapping of Short Grain Mutant Gene on Chromosome 5

#### 3.3.1. Mapping in an Expanded F_2_ Population

To further refine the genomic interval associated with the SG phenotype, the IC206-based F₂ population was expanded to 1,031 individuals, which segregated at a 3.5:1 WT:SG ratio, consistent with single-gene inheritance (*p* = 0.13). The slight deviation from a 3:1 ratio is consistent with sampling variation.

To further narrow the candidate region defined by recombination breakpoints, additional marker density was introduced within the interval. Eight PACE SNP markers spanning the 0.61–4.66 Mbp interval on chromosome 5 were used for genotyping, identifying 385 recombinant individuals, including 340 single recombinants and 45 double recombinants (Supplementary Table S4). To minimize ambiguity, only single recombinants were retained for subsequent mapping.

Recombinant haplotypes localized the SG locus to a ∼950 kb interval between SNP5 (3.00 Mbp) and SNP7 (3.95 Mbp (Fig 2a). SNP6 at 3.34 Mbp co-segregated perfectly with the SG phenotype in all informative recombinants and remained fully associated when genotyped across double recombinants. The presence of recombinants flanking SNP6 indicated that additional marker density in this region would further refine the locus.

**Fig 2:**
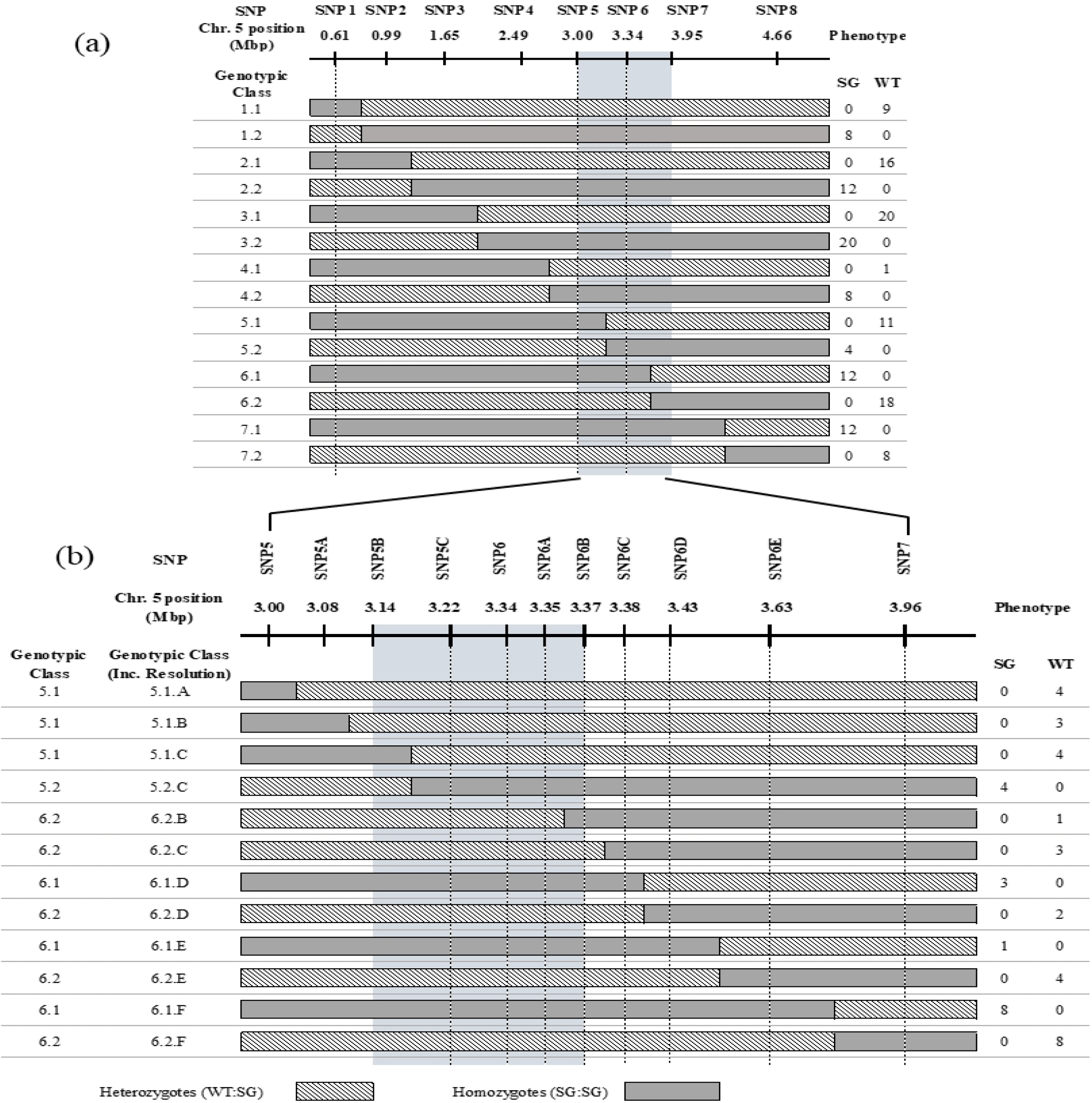
Mapping of the SG locus on Chr. 5 using F₂ recombinants. The x-axis shows physical position (Mbp); vertical bars indicate SNP positions. Each horizontal bar is a recombinant genotypic class. (a) Initial mapping within the 0.61–4.66 Mbp interval. (b) Higher resolution mapping within the 3.00–3.95 Mbp interval between SNP5 and SNP7; SNP5, SNP6, and SNP7 were re-genotyped for continuity. Gray = homozygous SG, hatched = heterozygous, white = homozygous WT. The light blue shaded region identifies the genomic interval containing the SG mutant gene.

Eight additional PACE markers were subsequently genotyped between SNP5 and SNP7, refining the interval to 233 kb between SNP5B (3.14 Mbp) and SNP6B (3.37 Mbp) (Fig 2b). Recombinant breakpoints on both sides of this interval collectively constrained the SG locus to this region. Complete recombinant and phenotypic dataset is provided in Supplementary S4 and S5 Tables.

#### 3.3.2. Progeny Testing for High-Resolution Fine-Mapping

To resolve the refined interval and further delimit the candidate region, F₃ progeny tests were conducted using selected F₂ recombinants. Within the 3.14–3.37 Mbp interval, F₂ recombinants were genotyped with 10 markers spanning this region. Because several recombinants were heterozygous or homozygous WT in the F₂ generation and therefore not phenotypically informative, F₃ progeny tests were conducted. Four previously excluded recombinants and one double recombinant were included, along with nine recombinants from earlier mapping.

Up to 24 F₃ individuals per family were evaluated, and segregation patterns were used to infer F₂ genotypes. All F₂ plants exhibiting the SG phenotype produced F₃ progeny fixed for SG, consistent with recessive inheritance. Informative recombinants collectively defined the proximal and distal boundaries of the SG locus, localizing it to a 41 kb interval between 3.19 and 3.23 Mbp on chromosome 5 (Fig 3; Supplementary Table S5). At this stage, all informative recombinants had been exhausted. The final 41 kb interval was defined by the nearest proximal and distal recombination breakpoints observed among the informative recombinant families, and no additional recombination events were available to further reduce the interval.

**Fig 3:**
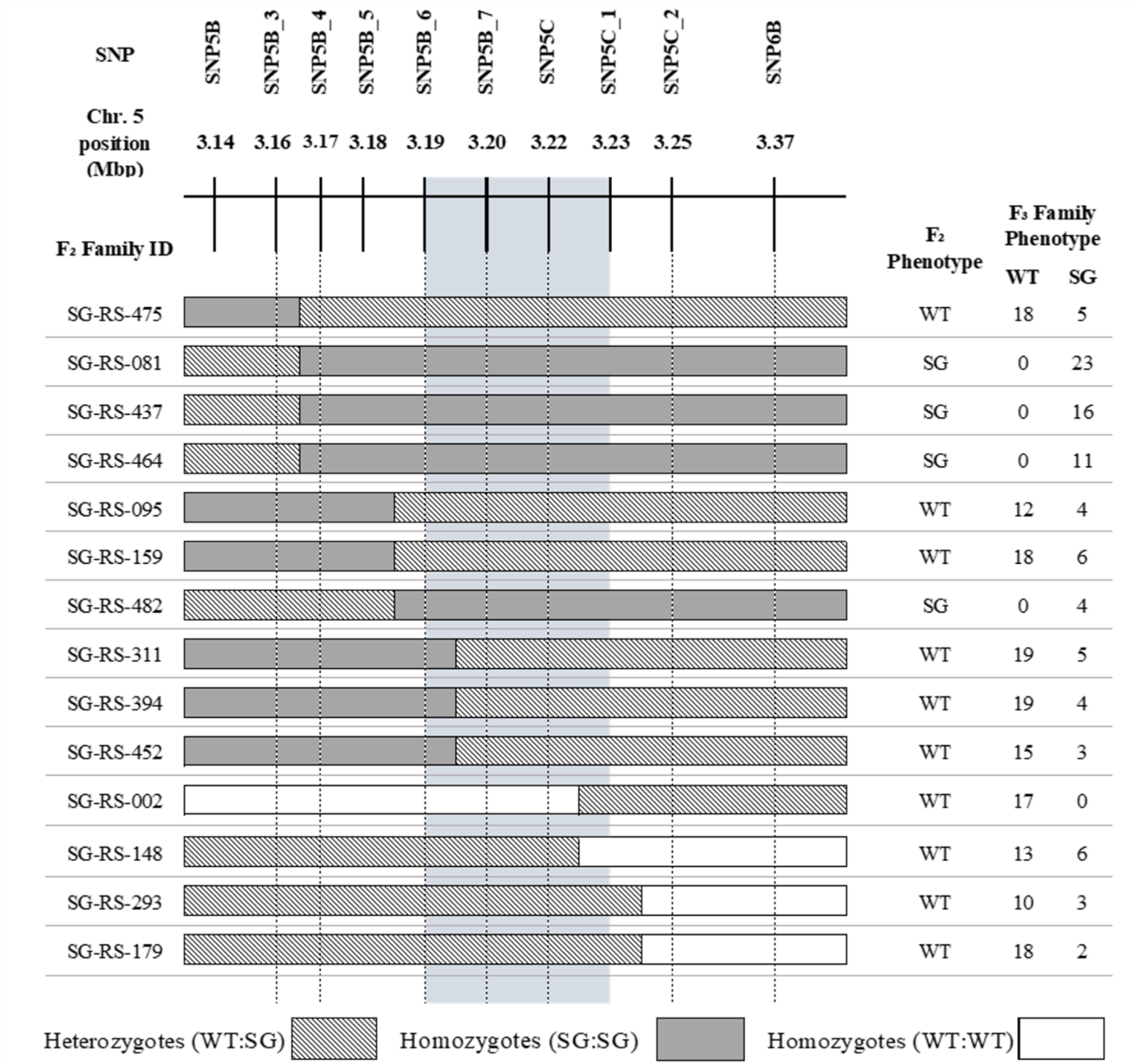
High-resolution fine-mapping using F₃ progeny from 14 recombinant F₂ families. The x-axis shows physical position (Mbp); vertical bars indicate SNP positions. Horizontal bars display the F₂ genotype of each F_3_ family. The phenotype of the F_2_ and the counts of each phenotype for the F_3_ families are shown to the right. Gray = homozygous SG, white = heterozygous, hatched = homozygous WT. The light blue shaded region identifies the refined mapped interval containing the SG locus.

### 3.4. Sequence Analysis and Identification of Candidate Causal Variant

Following fine-mapping to a 41 kb interval, whole-genome sequencing was used to identify candidate causal variants within the region. Annotation of the 41 kb fine-mapped interval identified five predicted genes, including the previously characterized grain-size gene *SMALL AND ROUND SEED 3* (*SRS3*), which encodes a kinesin-13 protein involved in cell elongation. None of the other annotated genes within the interval had known roles in grain size or morphology.

Whole-genome sequencing was performed on seven lines: WT RU2002174, WT IC206, the RU2002174-derived SG mutant, two F₂ individuals homozygous for the SG allele across the interval, and two F₂ individuals homozygous for the WT allele. Variant analysis focused on the 3.19–3.23 Mbp region of chromosome 5. Variants shared between WT RU2002174 and IC206 were excluded as background polymorphisms predating the mutation.

After filtering, only a single variant was present that distinguished all SG lines from WT lines: a G→T transversion at position 3,204,936 bp in exon 4 of *Os05g06280* (*SRS3*). This mutation introduces a premature stop codon (GAA→TAA), predicting a truncated 225-amino-acid protein (Fig 4). Although functional validation experiments (e.g., complementation or gene editing) were not performed and would be required to definitively confirm causality, the presence of this unique loss-of-function mutation, its complete co-segregation with phenotype, its location within the fine-mapped interval, and the strong phenotypic similarity to previously described *srs3* mutants collectively provide strong evidence that *SRS3* is the causal gene underlying the SG phenotype. The identification of a single loss-of-function SNP within *SRS3* is consistent with the observed recessive inheritance of the SG phenotype and provides a direct molecular explanation for the trait.

**Fig 4:**
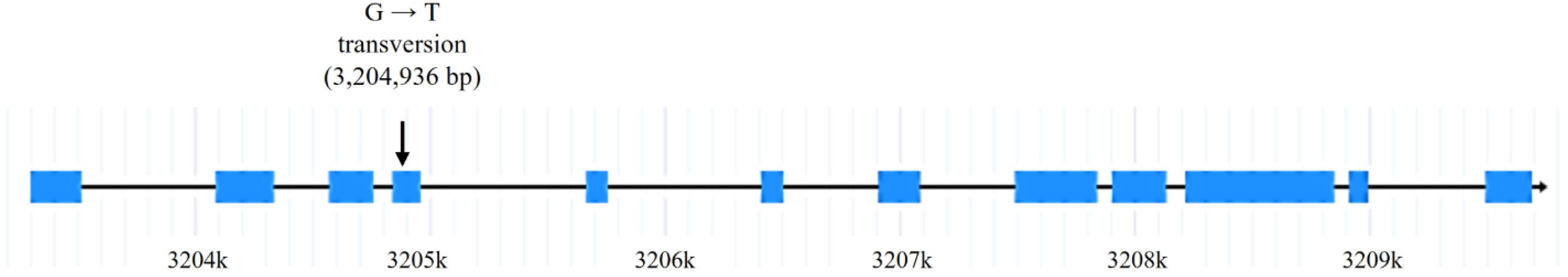
Candidate causal mutation identified in *SRS3* (Os05g06280). Sequence analysis of the 41-kb fine-mapped interval identified a single G→T transversion at position 3,204,936 bp in exon 4 of *SRS3*. The mutation converts a glutamic acid codon (GAA) to a premature stop codon (TAA), resulting in a predicted truncated protein. The schematic illustrates the exon-intron structure of *SRS3* and the position of the identified SNP. Adapted from RAP-DB [58].

### 3.5. Marker Development and Validation

A PACE assay targeting the exon 4 G→T polymorphism in *SRS3* was developed and used to genotype 251 F₃ individuals from recombinant families and 384 previously identified F₂ recombinants. In all cases, marker genotype showed complete concordance with grain phenotype, confirming the diagnostic utility of the marker.

To assess the novelty of the SG allele, the assay was applied to panel of 380 U.S. breeding lines (Native Trait Panel), and sequence data from approximately 3,000 global rice accessions were examined using the IRRI SNP-Seek database [59]. Only the WT allele was detected in all surveyed materials, indicating that the SG mutation in RU2002174 was not detected in representative U.S. breeding germplasm or in the approximately 3,000 accessions surveyed through SNPSeek, supporting its classification as a novel allele.

## 4. Discussion

### 4.1. Discovery of a Recessive Spontaneous Mutation and Breeding Program Implications

Spontaneous mutations represent a continuous but underexplored source of genetic variation in crop plants, particularly within advanced breeding populations where genetic uniformity is assumed. In this study, a short-grain (SG) off-type identified during routine seed increase was shown to be nearly identical to the wild-type (WT) line based on genome-wide SNP markers, effectively ruling out outcrossing as a likely explanation. In addition, the SG allele was absent from representative U.S. breeding germplasm and global rice diversity panels. Together, these observations strongly suggest that the SG phenotype did not arise from seed contamination or introgression. While alternative explanations cannot be completely excluded, the most parsimonious explanation is that this mutation arose de novo within the RU2002174 breeding line. RU2002174 was first evaluated in 2018, after which seed increase was maintained through hand-selected panicles. Based on the observed segregation pattern and advancement history of RU2002174, the mutation was inferred to have arisen during the 2017 seed increase cycle and remained heterozygous in 2018, when it would not have produced an SG phenotype. Some panicles selected for advancement in 2018 likely carried the SG allele in the heterozygous state, resulting in the panicle-row segregation for the SG phenotype first observed in 2019 during seed advancement.

The significance of this case lies not in the magnitude of the phenotypic effect alone, but in the documentation of a spontaneous single-gene mutation traced from initial field observation through genetic mapping and molecular identification. Such events are rarely characterized at this resolution despite their relevance to breeding operations. Because spontaneous mutations can compromise varietal uniformity, stability, and market classification, particularly when they affect major quality traits, this study reinforces the importance of continued monitoring of off-types during seed increase and purification, even in near-fixed materials [60, 61].

The SG phenotype clearly meets the criteria for short-grain rice, with reduced grain length and a length-to-width ratio below two. Although morphologically similar to short bold grains commonly associated with temperate *japonica* varieties favored in East Asian markets, this mutation arose in a tropical *japonica* genetic background characterized by high amylose content and intermediate gelatinization temperature. This combination of grain size and physicochemical properties is atypical, highlighting how spontaneous mutations can generate novel trait combinations that fall outside conventional market classes.

### 4.2. Characterization of the Spontaneous Mutation in the *SRS3* Gene Region

Together, the mapping, sequencing, and segregation data converge on a single highly supported candidate causal variant, linking the observed phenotype to a defined molecular lesion in *SRS3.* Fine-mapping localized the SG phenotype to a 41-kb interval on chromosome 5 containing *Os05g06280*, previously described as *SMALL AND ROUND SEED 3* (*SRS3*). Sequence analysis identified a single G→T transversion in exon 4 of *SRS3*, resulting in a premature stop codon and a predicted truncated protein. This mutation differs from previously reported induced *srs3* alleles, which were associated with mutations in later exons and identified through chemical mutagenesis [29, 30].

*SRS3* encodes a kinesin-13 protein involved in regulating cell elongation during seed development. Prior work demonstrated that *srs3* mutants exhibit shortened seeds due to reduced longitudinal cell length in lemma epidermal cells, implicating cytoskeletal dynamics as a key determinant of grain morphology. The truncation identified in this study is predicted to substantially disrupt protein function, consistent with the observed reductions in grain length of approximately 17–40% depending on grain type and the associated increase in grain width observed in the SG mutant. While the phenotypic outcome aligns with previous reports of *srs3* mutants, the discovery of a spontaneous loss-of-function allele in elite breeding germplasm provides additional independent support for the gene’s role and expands the allelic series at this locus.

Importantly, this work demonstrates that even well-characterized genes can acquire novel functional alleles through spontaneous mutation within breeding programs. Such alleles may go unnoticed or be eliminated during routine purification unless explicitly tracked and characterized. From a genetic perspective, the mapping strategy employed here illustrates how classical recombination-based approaches, combined with targeted sequencing, remain effective for resolving major-effect traits to gene level when population sizes and phenotyping are sufficient.

### 4.3. Broader Implications and Future directions

Previous studies of induced *srs3* mutants reported pleiotropic effects beyond grain morphology, including erect leaves, shortened internodes, and altered panicle architecture [30]. While the present study focused on grain size and genetic identification of the candidate causal mutation, traits such as leaf architecture, plant height, internode elongation, and panicle morphology were not systematically evaluated. Further work is needed to determine whether other pleiotropic effects are expressed in the RU2002174 SG mutant within this tropical *japonica* background. In particular, the extent to which reduced grain size may be compensated by changes in grain number or other yield components remains to be determined.

In addition, the unusual combination of short-grain morphology with high amylose content and intermediate gelatinization temperature presents an opportunity to further explore the relationship between grain shape and cooking quality. Although short-grain rice is often associated with low amylose and sticky texture, the SG mutant described here decouples grain size from starch properties, providing a useful genetic resource for studying how these traits interact to influence consumer perception and end-use quality.

## 5. Conclusion

This study documents the real-time emergence of a spontaneous, recessive mutation affecting grain size detected within breeding rows of rice. The mutation was mapped to a 41 kb interval on chromosome 5 and sequencing of the target region identified a single candidate causal nucleotide change consisting of a G→T transversion in exon 4 of Os05g06280 that introduces a premature stop codon in the *SRS3* gene. These findings report a novel mutation in a previously known gene and provide evidence of continued generation of naturally-occurring genetic variation even in advanced breeding materials and highlight the importance of vigilant monitoring of off-types due to their potential impacts on varietal uniformity, trait stability, and market suitability.

## Acknowledgements

The authors thank Jennifer Manangkil for assistance with phenotypic evaluations and Jennifer Dartez and Madeline Lejeune for assistance with genotyping activities. The authors also thank the staff at the H. Rouse Caffey Rice Research Station for contributions to this work. Funding support was contributed in part by the Louisiana Rice Research Board.

## Author Contributions

Conceptualization: A.F., M.G.M., B.A. Data curation: M.G.M. Formal analysis: M.G.M., A.F. Funding acquisition: A.F. Investigation: M.G.M., B.A., J.K.R., A.F. Methodology: M.G.M., B.A., J.K.R., A.F. Writing – original draft: M.G.M., A.F. Writing – review & editing: all authors

## Competing interests

The authors have declared that no competing interests exist.

## Data Availability

All sequencing data are available in the NCBI Sequence Read Archive (SRA) under BioProject PRJNA1472089. All other relevant data are provided within the manuscript and its Supporting Information files.

## Supplementary Information

**S1 Table:** Mean grain measurements for WT and SG phenotypes of RU2002174, including grain length (L), width (W), thickness (T), and length-to-width ratio (L/W) for rough rice, brown rice, and milled rice.

**S2 Table:** Details of PACE SNP markers, including SNP number (SNP #), LSU marker identification (LSU ID), chromosome (Chr.), physical position (bp), and primer sequences (Allele 1, Allele 2, and common primer).

**S3 Table:** Genotype data for chromosome 5 SNP markers significantly associated with the SG phenotype in the LSU500 analysis of the IC206 × RU2002174 F₂ population. Marker information includes SNP number (SNP #), LSU marker identification (LSU ID), chromosome (Chr.), physical position (bp), and parental alleles. Phenotypes are classified as WT (long grain) or SG (short grain).

**S4 Table:** Rough mapping on Chromosome 5 using 8 SNP markers, including recombinant classes identified among 340 single recombinants and 45 double recombinants in the F₂ population. Phenotypes are classified as WT (long grain) or SG (short grain), with SG** indicating variable grain size within a panicle. Columns include line identification (Line ID), Population (Pop 1, refers to population 1; Pop 2, refers to Population 2), recombinant class (Rec. Class), genotypic class (Gen. Class), and phenotype class (Phenotype). It also shows SNP Number (SNP #), LSU identification for each marker (LSU ID), Chromosome (Chr.), and SNP position (Position (bp)). Additionally, allele 1 and allele 2 are shown for each SNP marker, detailing the corresponding nucleotide base and the associated WT and SG parental haplotypes.

**S5 Table:** Increased resolution mapping data using 9 additional SNP markers together with SNP5, SNP6, and SNP7 across 45 single recombinant F₂ plants. The table includes recombinant F₂ lines, corresponding F₃ progeny test results, genotype classifications, and marker information used to delimit the SG locus to the final mapped interval. Phenotypes are classified as WT (long grain) or SG (short grain).

## Notes

### Competing Interest Statement

The authors have declared no competing interest.

